# Development of genomic resources for cattails (*Typha*), a globally important macrophyte genus

**DOI:** 10.1101/2023.04.21.537876

**Authors:** Alberto Aleman, Marcel E. Dorken, Aaron B. A. Shafer, Tulsi Patel, Polina A. Volkova, Joanna R. Freeland

## Abstract

**1. Background:** A critical knowledge gap in freshwater plants research is the lack of genetic tools necessary to answer fundamental questions about their demographic histories, adaptation, and phylogenetic relationships. One example of this is *Typha*, a global genus of freshwater plants foundational to wetlands that is also becoming an increasingly problematic biological invader in numerous regions worldwide; while important insights have been discovered for this genus, existing markers are insufficient to answer fundamental questions about their demographic histories, adaptation, and phylogenetic relationships, to identify introduced and hybrid lineages, and to examine patterns of hybridisation and introgression.

**2. Methods:** We optimised a library preparation and data processing protocol to develop genome–wide nuclear and plastid resources for studying the evolutionary history, genetic structure and diversity, hybridisation, local adaptation, invasiveness, and geographic expansion dynamics of *Typha*.

**3. Main results:** We sequenced 140 *Typha* samples and identified ∼120K nuclear SNPs that differentiate *T. angustifolia*, *T. domingensis* and *T. latifolia* and retrieved their plastome sequences. We observed genetic introgression among the three species.

**4. Conclusions:** Following a fast, straightforward, and cost–efficient genomic library preparation protocol, we produced a suite of genome–wide resources to facilitate investigations into the taxonomy and population genetics of *Typha* and to advance the genomic understanding of wetland plants.

**5. Contributions:** The protocol described, the updated chromosome–level genome assembly of *T. latifolia*, the catalogue of species-specific SNPs, and the chloroplast sequences produced in this study comprise permanent resources that can be applied to study the genetic composition of multiple populations and hybrid zones and will be incorporated into future studies of *Typha,* an ecologically important and globally invasive macrophyte.

## Introduction

Freshwater plants are essential to aquatic ecosystems, shaping their habitats’ structure and ecological functions (Chambers et al., 2008; Christie et al., 2009; Rejmankova, 2011). Although freshwater plants have been increasingly incorporated into applications that include habitat restoration and invasive species management, they remain highly understudied compared to terrestrial plants (Evangelista et al., 2014; Iversen et al., 2022). One key knowledge gap in freshwater plants research is the lack of genetic tools necessary to answer fundamental questions about their demographic histories, adaptation, and phylogenetic relationships (Fay et al., 2019; Maréchal, 2019; O’Hare et al., 2018; Yannelli et al., 2022).

Genetic characterisation of freshwater plants has been hampered by biological and technical challenges, plus biases in scientific research (Matheson & McGaughran, 2022; Troudet et al., 2017). Hundreds to thousands of molecular markers are often required for genomic–based research on topics such as gene flow and adaptation of non–model organisms (da Fonseca et al., 2016; Stapley et al., 2010), but the *de novo* development of genomic resources can be both time–consuming and expensive (Hu et al., 2020; Ortega et al., 2020; Prieto et al., 2021). Overcoming these challenges is now feasible using novel, rapid and cost–effective methods that capture genome–wide genetic variation, allowing researchers to address questions related to taxonomy, evolution, and conservation (Andrews et al., 2016; Goodwin et al., 2016).

*Typha* L. (cattails) is a global genus of rhizomatous perennial, monoecious, self-compatible, and wind-pollinated freshwater plants foundational to wetlands (reviewed in Bansal et al., 2019). Cattails are a valuable ecosystem resource and play a fundamental ecological role by cycling nutrients, preventing erosion, maintaining stable water levels and providing food and shelter for wildlife (Andrews & Pratt, 1978; Bonanno & Cirelli, 2017; Dieye et al., 2017; Kimmerer, 2013; Svedarsky et al., 2019). One major challenge in *Typha* research has been taxonomic identification, which cannot be fully accomplished using morphological characters due to their high intraspecific variability and interspecific hybridisation. Consequently, the richness of cattail species, their taxonomy, provenance (i.e., alien or native lineages), and phylogenetic relationships remain unsolved (Ciotir & Freeland, 2016; Volkova & Bobrov, 2022; Zhou et al., 2018). A refined *Typha* taxonomy along with species-specific genetic markers are necessary to identify introduced and hybrid lineages, which are increasingly documented as invasive, e.g., *T. domingensis* Pers. in Central America, *T.* × *glauca* Godr. (*T. angustifolia* L. *× T. latifolia* L.) in North America, and *T. latifolia* L. in Oceania and Western Europe, (Bansal et al., 2019; *GISD*, n.d.; Govaerts, 2004; Hall, 2009; Maldonado, 2019; Xu et al., 2013).

High–throughput sequencing technologies, novel, cost and time–accessible genome library preparations, and the recent assembly of the *T. latifolia* genome (287.19 Mb) (Goodwin et al., 2016; Rowan et al., 2019; Widanagama et al., 2022) collectively present an opportunity to develop a suite of genomic resources for *Typha*. In addition to taxonomic resolution, these resources will facilitate investigations of the evolutionary history, genetic structure and diversity, hybridisation and introgression, local adaptation, invasiveness and geographical expansion of this genus. We applied a high–throughput sequencing protocol for enzymatic fragmentation, library preparation, and data processing and produced genome–wide resources for three *Typha* spp. By optimising the method from Rowan et al. (2019), we generated a catalogue of nuclear SNPs that characterise *T. angustifolia*, *T. domingensis*, and *T. latifolia* that include chloroplast–genome sequences in a fast, straightforward, and cost–efficient manner.

## Materials and methods

### Reference genome

We used the reference–based scaffolder Chromosomer 0.1.4a (Tamazian et al., 2016) to align the 1158 *T. latifolia* scaffolds from Widanagama et al. (2022) (Genbank accession: JAIOKV000000000.1) with the *T. latifolia* isolate L0001 (15 chromosomes, GenBank accession: JAAWWQ000000000.1) and produce a local chromosome–level *T. latifolia* genome. Widanagama et al. (2022) had a significantly higher mapping success of unrelated re–sequenced *Typha* spp. compared to the isolate L0001, suggesting it is a more representative *Typha* genome assembly. The scaffolds were aligned as chromosomes using BLAST+ 2.12.0 (Camacho et al., 2009) with the software default settings, and the alignments were anchored in Chromosomer, establishing a gap length = 0 and a ratio threshold = 1.

### Sampling, DNA extraction, and sequencing

Samples were either obtained from previous studies or collected across Eurasia, and DNA was extracted at Trent University following published protocols (Bhargav et al., 2022; Ciotir et al., 2017; Pieper et al., 2020, 2017; Tangen et al., 2022; Tisshaw et al., 2020). Briefly, leaf tissue was dried in desiccant silica beads and stored at −20°C. Dried leaf material was ground with a Retsch® MM300 mixer mill (Haan, Germany). DNA was extracted from 25 - 30 mg of semi-fine powder of each sample using the EZNA Plant DNA kit (Omega-Bio Tek) or the Fastpure plant DNA isolation mini kit (Nanjing Vazyme Biotech-China) protocols for dried material, with a final elution of 100 μL. We obtained DNA from 38 *T. angustifolia*, 25 *T. domingensis*, and 77 *T. latifolia* samples (n = 140) (Figure 1; Supplementary Table S1) previously identified to taxon using a combination of genetic analyses of microsatellite loci (Kirk et al., 2011; Snow et al., 2010) and morphological characteristics (Grace & Harrison, 1986; Smith, 1967). Extracted DNA was quantified using a Qubit fluorometer (Thermofisher Scientific) and calculated as the mean of three independent readings for each sample. All samples were either standardised to 2 ng/μL by dilution with nuclease–free water or left undiluted if at concentrations less than 2 ng/μL (0.4 –1.9 ng/μL).

**Figure 1.**
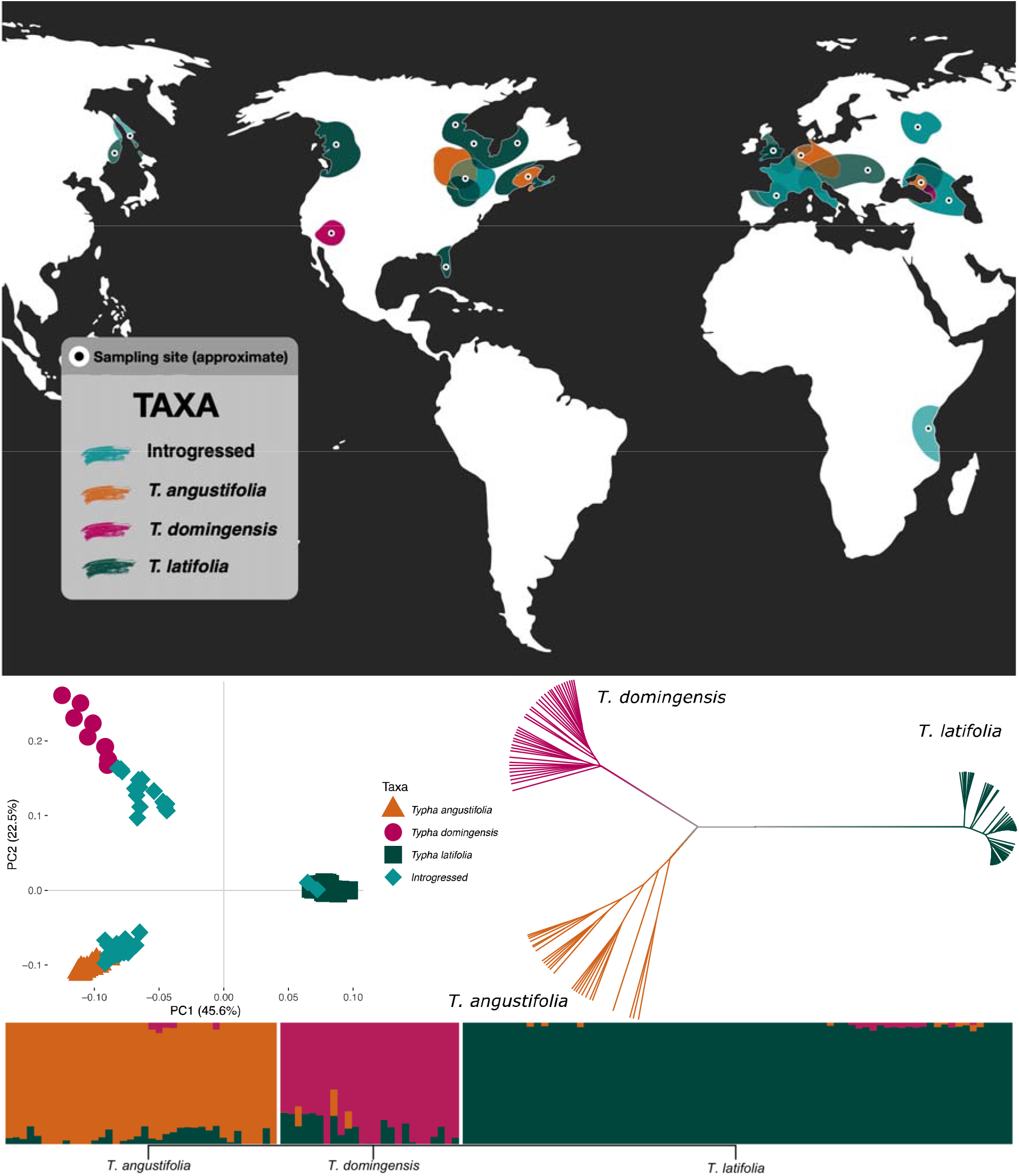
Genetic structure results of 12,177,703 nuclear SNPs obtained for three *Typha* spp. Top: Sampling locations in this study. Black points indicate approximate sampling sites, and coloured areas indicate the taxa identified. The number of samples is not shown. Left: Principal Component Analysis for PC1 and PC2. Shapes represent individuals, and colours represent taxa, as in the box. Right: Neighbour–joining tree. Branches represent individuals, and colours indicate species belonging as labelled. Bottom: ADMIXTURE (*K* = 3). Vertical bars represent individuals, and the admixture proportion is shown with different colours.

Rowan et al. (2019) reported a relatively rapid and cost-effective library preparation technique for genomic sequencing by enzymatic fragmentation followed by ligation of short adapter sequences (i.e., tagmentation) using transposases (Nextera XT) and relatively low yields of DNA. Our protocol was based on this method with a few modifications. First, each DNA sample was tagmented with the Illumina Tagment DNA enzyme (TD) and buffer kit (small kit, #20034210). As the ratio of TD enzyme to DNA is crucial for the reaction, we initially followed Rowan et al. (2019) and subsequently optimised the reagent volumes for our library preparation as 5.5 μL of 5*×* TD buffer, 0.5 μL of 1*×* TD enzyme and 4 μL of DNA (standardised or undiluted), keeping all reagents on ice during the preparation. Samples were incubated at 55°C for 10 minutes and left at room temperature for five minutes, and 5 μL of each sample were run on an agarose gel to confirm the efficacy of the tagmentation reaction, evidenced by visible smears. The tagmented DNA was then amplified using unique dual indexing based on combinations from a total of 24 N7 (47 bases) and 8 S5 (51 bases) adapters (Alpha DNA, Canada). The PCR cocktail included 0.2 µM of each index, 0.5U of KAPA HiFi HotStart DNA polymerase (Roche), 12.5 μL of 5*×* KAPA reagent, 5 μL of tagmented DNA, and 6.5 μL of nuclease–free water to a final volume of 25 μL. The PCR cycle comprised 72°C (3 minutes); 95°C (30 seconds); and 14 cycles of 95°C (10 seconds), 55°C (30 seconds), and 72°C (30 seconds). Once again, visible smears confirmed amplification success after running 5 μL of the PCR product on an agarose gel; then, 10 μL of each sample were pooled, and the remaining PCR products were stored at −20°C. The pooled library was purified with a QIAquick PCR purification kit (QIAGEN) following the manufacturer’s protocol, with a finalelution in 50 μL of elution buffer. The library was quantified using a D1000 Tapestation assay (Agilent Technologies, USA) and a Qubit fluorometer (Thermo Fisher Scientific). A quality–control paired–end sequencing was executed using a Miseq (151 bp) to ensure the genomic library was compiled successfully. Finally, paired–end sequencing was performed on a Novaseq 6000 (126 bp) at The Centre for Applied Genomics (Toronto, Ontario).

### Raw data processing, filtering, and SNP–calling

The quality of the demultiplexed raw sequences was evaluated using FastQC 0.11.9 (Andrews, 2017) and MultiQC 1.14 (Ewels et al., 2016). Read pairing and adapter pruning were carried out with trimmomatic 0.39 (Bolger et al., 2014), removing any cleaned reads shorter than 100 bp. Paired and remaining unpaired reads were mapped to our chromosome–level *T. latifolia* nuclear genome plus the *T. latifolia* plastome reference (Genbank accession: NC_013823.1) using the mem module of BWA 0.7.17 (Li & Durbin, 2009). Mapped reads from Miseq and Novaseq 6000 sequencers were merged, and mapping statistics were evaluated with the flagstat and coverage modules of SAMtools 1.15.1 (Li et al., 2009).

Genotype-calling was performed with ANGSD 0.93 (Korneliussen et al., 2014) following the SAMtools model, retrieving hard-called SNPs with a minimum *p*–value of 1e^−6^, minimum mapping and sequencing qualities of 20, discarding any indels and triallelic sites, and outputting a binary Variant Call Format file (*-doGeno 4 -gl 1 -skipTriallelic 1 -SNP_pval 1e-6 -minMapQ 20 -minQ 20 -doMajorMinor 1 -domaf 1 -doPost 1 -* doBcf 1).

For the nuclear analyses, SNPs with > 50% missing data across all samples and sites mapped to the plastome were removed in VCFtools 0.1.16 (Danecek et al., 2011). We did not apply any additional filters to SNP identification: our samples represent a broad geographical sampling (Figure 1; Supplementary Table S1) and thus were not expected to be in Hardy–Weinberg equilibrium; additionally, as allele frequencies were unlikely to be representative of regional allele frequencies, we did not apply a minor allele frequency filter; neither did we filter for linkage equilibrium, as eliminating alleles that are in linkage disequilibrium is likely to decrease the resolution to detect hybridisation and introgression (Alexander, 2020; Pearman et al., 2022).

### Genetic structure and diagnostic markers

We used nuclear SNPs to assess the most likely number of genetic clusters across all samples and the membership of each plant to these clusters using three complementary approaches: i) ADMIXTURE 1.3.0 (Alexander & Lange, 2011) was run with *K* = 1 – 10, and the optimal number of clusters was chosen via the cross–validation procedure, ii) a neighbour–joining tree from the samples’ pairwise genetic distance matrix (expressed as allele counts, transformed on the R 4.2.1 package ape 5.7-1 (Paradis et al., 2004; R Core Team, 2022)), and iii) a Principal Component Analysis (PCA) were performed with Plink 1.90 (Purcell et al., 2007). To avoid over- or under-estimating genetic structure (Janes et al., 2017), we verified that the assignment of samples to genetic clusters (corresponding to three species, see *Results*) was consistent for each approach.

Potential introgression was tested by running ADMIXTURE (*K* = 1 – 5) on three datasets, each comprising a combination of two genetic clusters, using only those SNPs that remained variable based on the two species being compared. We confirmed that the cross–validation procedure for the runs of each species’ pair chose the optimal number of clusters as two (*K* = 2) and used the admixture proportion (*Q score*) as an index of potential introgression of each sample. Applying Senn & Pemberton (2009) and Smith et al. (2018) thresholds, individuals whose *Q* score was 0.05 ≤ *Q* ≤ 0.95, were considered as genetically introgressed.

To compare the levels of differentiation between clusters, values of Weir and Cockerham’s genetic differentiation (*F_ST_*; Weir & Cockerham, 1984) and genetic divergence (*d_xy_*; Nei & Miller, 1990) for every variable site in 10 Kbp windows between species pairs were computed with pixy 1.2.7 (Korunes & Samuk, 2021), and the means were calculated. Species–specific SNPs were identified i) for each species’ pair and ii) by running three paired comparisons of one species versus the other two on each run, using DiagnoSNPs 1.0 (Arce-Valdés, 2022). We removed genetically introgressed individuals before estimating levels of genetic differentiation and identifying species–specific SNPs.

### Chloroplast genome reconstruction and phylogenetic analysis

We implemented a reference–guided workflow to reconstruct whole–chloroplast–genome sequences. Nucleotide calling was performed individually for each of the 140 samples in ANGSD, using the reads that mapped to the plastome reference, requiring minimum mapping and base qualities of 20, and using N’s for missing data (*-dofasta 2 - minMapQ 20 -minQ 20 -doCounts 1*). The sequences were aligned to the chloroplast genomes of *T. przewalskii* Skvortsov, *T. lugdunensis* P. Chabert, *T. orientalis* C. Presl, and *Sparganium natans* (GenBank accessions: NC_061354.1, NC_061353.1, NC_050678.1, and NC_058577.1), following Smith et al. (2021) by applying MAFFT 7.0 default settings (i.e., with the flag *--nzero*) (Katoh et al., 2019). Snp–sites 2.5.1 (Page et al., 2016) was used to remove regions of the genome with ambiguous positions, gaps, and missing data, such that if any of those was found in a sequence, that position was removed for all sequences. Nucleotide diversity (π) was calculated in the R package pegas (Paradis, 2010).

The phylogenetic relationships of the chloroplast genome sequences were reconstructed in RAxML-NG 1.1 (Kozlov et al., 2019). Model selection was based on jmodeltest 2.1.10 results (Darriba et al., 2012) using the Akaike Information Criterion. RAxML was run under a TVM + G4 + I model with the automatic and thorough bootstrap options, starting from 100 random trees and employing *Sparganium natans* as the outgroup. The best–scoring ML tree was visualized.

## Results

### Genome scaffolding, mapping statistics, and genotyping

Approximately 99.81% of the scaffold sequences were aligned to the template genome. The scaffolds were anchored to 15 chromosomes producing a genome of 285.11 Mb (GenBank accession: JAIOKV000000000.2). The total assembled size was comparable to the *T. latifolia* genome sizes of Widanagama et al. (2022) (287.19 Mb) and the isolate L0001 (214.13 Mb). Updating the chromosome–level genome assembly simplified our downstream analyses while keeping the highest mapping success of unrelated re–sequenced *Typha* spp., and facilitating an accurate recombination map for future studies of speciation, hybridisation, and the genomic landscape of introgression in *Typha*.

After quality control, 982 M clean paired–end reads were retained, and ∼98% mapped to the reference genome. With minimum mapping and sequencing qualities = 20, the average depth and breadth of coverage for the nuclear sequences were 4× and 42%, respectively. Over 60% of the plastome breadth was covered across all samples (mean depth = 711×), enabling us to use 96,591 bp for the phylogenetic reconstruction. We assembled 12,177,703 bi–allelic nuclear SNPs across the 140 *Typha* samples (7,122,151 with a MAF > 0.05). The total genotyping rate, i.e., the mean proportion of samples with data for each SNP, was 0.68.

### Genetic structure and diagnostic markers

The admixture analysis, the PCA, and the neighbour–joining tree each established the most likely number of genetic clusters as three (*K* = 3) (Figure 1): in line with previous taxonomic identifications, 38 samples were identified within the *T. angustifolia* cluster (15 of which had *T. latifolia* introgression and 5 of which had both *T. domingensis* and *T. latifolia* introgression); 25 samples were in the *T. domingensis* cluster (one with *T. angustifolia* introgression, 12 with *T. latifolia* introgression, and 3 with both *T. angustifolia* and *T. latifolia* introgression); and 77 samples were in the *T. latifolia* cluster (one with both *T. angustifolia* and *T. domingensis* introgression, and one with *T. angustifolia* introgression). Using only the 18 *T. angustifolia*, 9 *T. domingensis*, and 75 *T. latifolia* non-introgressed samples, the mean pairwise interspecific *F_ST_* and *d_xy_* values ranged from 0.25 to 0.49 and 0.28 to 0.35 (Table 1), respectively, with *T. latifolia* showing the highest differentiation with both *T. angustifolia* and *T. domingensis.* We identified 119,324 nuclear species-specific SNPs by pairwise comparisons between the three species and 16,856 SNPs when one species was compared to the other two (*T. angustifolia* = 10,537; *T. domingensis* = 3,838; *T. latifolia* = 3,044).

**Table 1.** Mean Weir and Cockerham’s pairwise genetic differentiation (*F_ST_*) and genetic divergence (*d_XY_*) measured in 10 Kbp windows for every variable site, number of SNPs, and diagnostic markers (SNPs with fixed opposite alleles) between three *Typha* spp.

| Pairwise comparison | $F_{ST}$ | $d_{XY}$ | SNPs | Diagnostic SNPs |
| --- | --- | --- | --- | --- |
| *T. angustifolia* – *T. domingensis* | 0.25 | 0.28 | 10,358,977 | 33,436 |
| *T. angustifolia* – *T. latifolia* | 0.44 | 0.35 | 8,786,870 | 30,113 |
| *T. domingensis* – *T. latifolia* | 0.49 | 0.34 | 9,380,607 | 55,775 |

### Chloroplast genome reconstruction and phylogenetic relationships

After removing all ambiguities and missing data from the chloroplast genomes, we were left with an alignment of 96,591 bp across all 140 sequences and the four references, with 4,916 segregating sites and π = 0.003. The phylogenetic reconstruction was congruent with the nuclear genetic structure results (i.e., individuals were consistently assigned to the same nuclear and plastid lineages), grouping the 143 *Typha* samples into three lineages, with *T. angustifolia* in one clade, *T. domingensis* and *T. orientalis* sharing another, and *T. latifolia* and *T. przewalskii* in a third one (Figure 2). All interspecific nodes were strongly supported.

**Figure 2.**
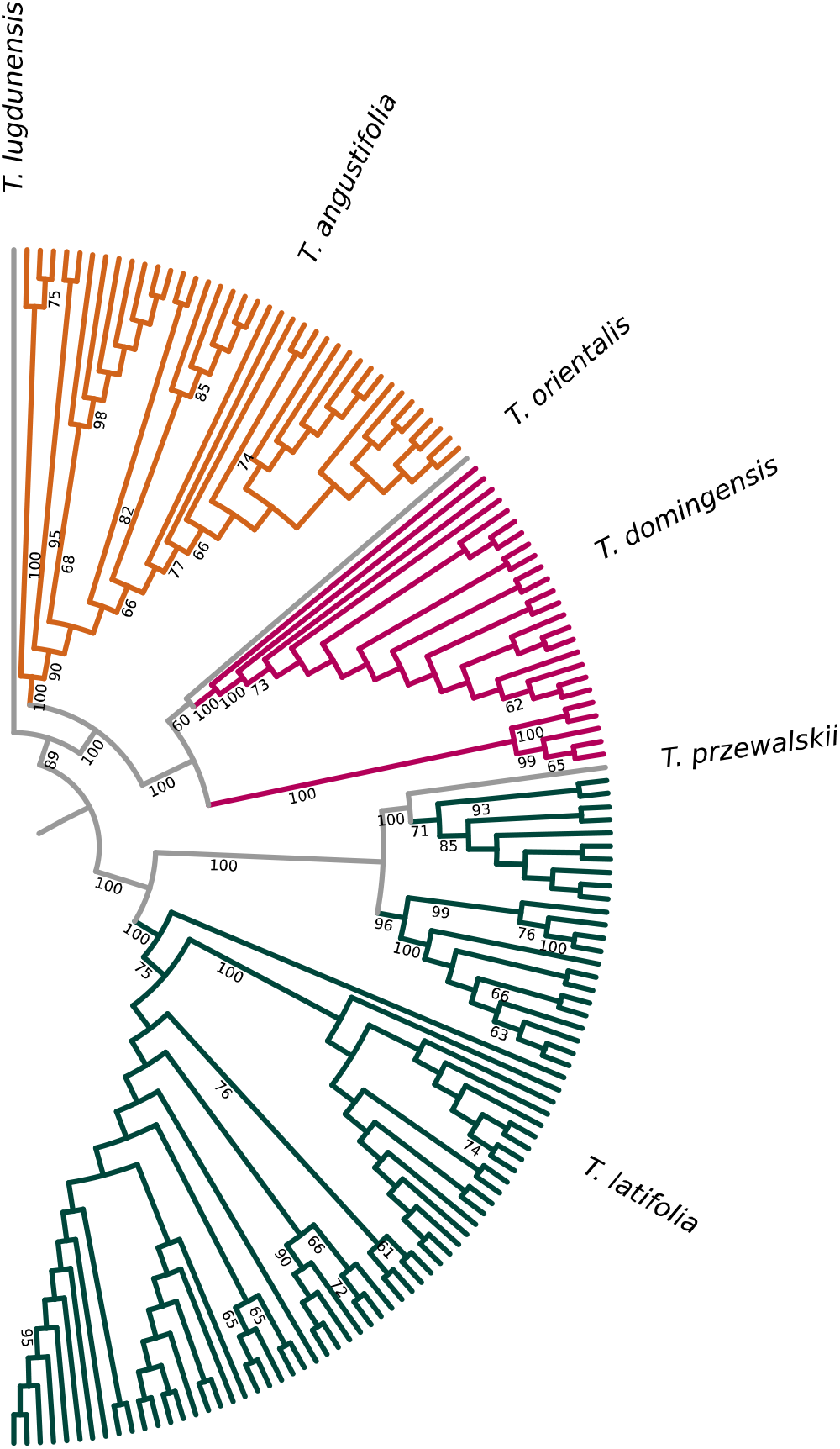
Chloroplast phylogeny of 140 samples from three *Typha* spp. and four NCBI references, based on 96,591 bp. The tree was produced with *Sparganium natans* as the outgroup and drawn without it. Branches represent individuals, and colours indicate species belonging as labelled. Numbers indicate branch support ≥ 60. Branch lengths are not shown.

## Discussion

We aimed to produce a suite of genome–wide resources to facilitate investigations into the taxonomy and population genetics of *Typha* and to advance the genomic understanding of wetland plants. Following a fast, straightforward, and cost–efficient genomic library preparation protocol (Rowan et al., 2019), we sequenced 140 *Typha* samples, obtaining an average breadth of 42% of the nuclear genome, characterising 119,324 nuclear SNPs that collectively differentiate three *Typha* spp., and producing chloroplast sequences with a breadth of coverage > 60% per sample. With a cost below 15 USD per sample and a processing time of two hours for the library preparation, our workflow is a rapid and cost–effective protocol that can be applied in population genomic studies for investigating levels of genetic diversity and differentiation, identifying conservation units and alien taxa, and investigating hybrid zones, among other purposes. Additionally, our results reveal the feasibility of reconstructing whole-chloroplast-genome sequences as a by-product of the enzymatic fragmentation for high-throughput sequencing libraries, making plastome research simpler and inexpensive for species with an available reference.

Three genetic clusters were identified from both nuclear and chloroplast genomes, corresponding to *T. angustifolia*, *T. domingensis*, and *T. latifolia*, and genetic introgression was detected among the three species. While there are some reports of hybridisation between *T. domingensis* and either *T. angustifolia* or *T. latifolia* (Ciotir et al., 2017; Govaerts, 2004; Smith, 1967), range-wide surveys are lacking. Future research should address the extent to which hybridisation is shaping the genetic differentiation and diversity of these three species. Furthermore, *T. domingensis* is increasingly invading regions in Nigeria (Ringim et al., 2016), Costa Rica (Trama et al., 2017) and North America, potentially expanding its range across the latter (Spencer & Vincent, 2013; Zhang et al., 2008). However, the taxonomic identity of these plants is unclear –are they hybrids, non–native lineages, native lineages responding to environmental change, or misidentified *T. angustifolia* (Bansal et al., 2019)? By characterising SNPs that differentiate *T. domingensis*, we provide a valuable resource to answer this and other questions across the evolutionary and hybridisation history of *Typha*.

The markers that differentiate *T. angustifolia* from *T. latifolia* will have important applications in North America, where the two species interbreed across a large area and produce an invasive interspecific hybrid (*T.* × *glauca*) that dominates wetlands, alters nutrient cycling, and reduces biodiversity across the Great Lakes Region (Bansal et al., 2019); additionally, this hybrid is expanding throughout the Prairie Pothole Region, causing native plant diversity to decrease in invaded potholes (Jones et al., 2023), and may impact essential habitat for millions of breeding and migratory waterfowl species (Tangen et al., 2022). Until now, molecular resources to characterise *T. angustifolia*, *T. latifolia*, and *T.* × *glauca* were limited to sets of relatively few individual markers that have produced important insights: RAPDs, chloroplast DNA sequences, and codominant SSR loci have contributed to exposing the sexual fertility of first–generation hybrids (F1s) (Snow et al., 2010), asymmetric hybridisation, with *T. angustifolia* being mainly the maternal parent (Ball & Freeland, 2013; Kuehn et al., 1999; Pieper et al., 2017), overall comparable levels of sexual and clonal reproduction in parents and F1s (Pieper et al., 2020; Travis et al., 2011), heterosis in F1s (Bunbury-Blanchette et al., 2015; Travis et al., 2010; Zapfe & Freeland, 2015), a high frequency of F1s in natural populations (Kirk et al., 2011; Travis et al., 2010), the capability of F1s to backcross, plus partial sterility in F1s coupled with hybrid breakdown of F2s and advanced–generation hybrids (Bhargav et al., 2022; Pieper et al., 2017). However, critical inquiries remain unresolved because existing markers are insufficient to expose the prevalence of advanced–generation hybrids and backcrosses in wild populations. The expansive suite of SNPs identified in this study will facilitate investigations into the extent of hybridisation, hybrid breakdown dynamics, and adaptive introgression across the *T. × glauca* hybrid zones, allowing researchers to understand the processes shaping this genus speciation and species boundaries, and to inform conservation and management strategies.

Fundamental genetic tools are essential for investigating freshwater plants’ biology, management, and conservation (O’Hare et al., 2018). The protocol described in this paper, the updated chromosome–level genome assembly of *T. latifolia*, the catalogue of species-specific SNPs, and the chloroplast sequences produced for each sample comprise permanent resources that can be applied to study the genetic composition of multiple populations and hybrid zones. Genome–wide sequencing techniques and reference–based chloroplast genome assemblies are promising tools to clarify the demographic histories, dispersal, adaptation, and taxonomy of multiple congeneric macrophyte species (Russello et al., 2015; Straub et al., 2012), and substantial genome–wide research will allow us to tackle these and other knowledge gaps in *Typha* and other freshwater taxa.

## Acknowledgements

We acknowledge that the laboratory procedures and data analyses were conducted at Trent University, which is on the traditional territory of the Mississauga Anishinaabeg. The Natural Sciences and Engineering Research Council of Canada (NSERC) financially supported this work, and Alberto Aleman is funded by the Environmental & Life Sciences Graduate Program at Trent University. The work of Polina A. Volkova was supported by the Russian Science Foundation grant no. 23-14-00115. We thank V. Bhargav, N. Tikhomirov, and M. Ivanova for providing plant tissue samples; T. Pimenov, M. Aksyonova and the staff of the Dagestansky Nature Reserve, in particular, G. S. Dzhamirzoyev, for their help in the field, AO “IEPI” for organising fieldwork in Krasnodar Region, and SHARCNET and Compute Canada for providing computational resources. Finally, we thank Camille Kessler for her comments on the manuscript and Enrique Ruiz for his work in Figure 2 (top).

## Author Contributions

Conceptualisation: ABAS, JRF, and MED. Developing methods, conducting the research, data interpretation and writing: AA, ABAS, JRF, MED, PAV, and TP. Data analysis and preparation figures & tables: AA.

## Data Accessibility Statement

The updated nuclear genome assembly will be submitted to GenBank (JAIOKV000000000.2). Code, species-specific SNP locations (chromosome, position) and chloroplast-genome sequences produced in this study will be available at https://gitlab.com/WiDGeT_TrentU/graduate_theses/-/tree/master/aleman/fwb and https://github.com/al-aleman/totoras_fwb. High-throughput sequencing raw data and Variant Call Format files are available from the authors upon reasonable request.

## Conflict of Interest

The authors declare no conflict of interest.

---

**Supplementary Table S1.** Sampling identifiers, geographic coordinates, species membership, and genetic introgression status of the 140 *Typha* collected for this study.

| Sampling ID | Latitude<br>(approximate) | Longitude<br>(approximate) | Species | Genetic<br>introgression |
| --- | --- | --- | --- | --- |
| BHP1 | 50.79 | -119.21 | *T. latifolia* | No |
| BJL2 | 50.57 | -119.61 | *T. latifolia* | No |
| BJL3_1 | 50.57 | -119.61 | *T. latifolia* | No |
| BRT2B | 44.74 | -63.24 | *T. angustifolia* | No |
| BUN1 | 53.89 | -122.81 | *T. latifolia* | No |
| C05TL | 52.72 | -1.37 | *T. latifolia* | No |
| CA13_TL | 44.85 | 7.71 | *T. latifolia* | No |
| CA3TL | 47.68 | 22.46 | *T. latifolia* | No |
| CC1_TL | 50.77 | 2.31 | *T. latifolia* | No |
| CL10TA | 50.77 | 2.31 | *T. angustifolia* | Yes |
| COTL6 | 52.72 | -1.37 | *T. latifolia* | No |
| CY6ATA | 51.43 | -0.11 | *T. angustifolia* | No |
| CY7ATA | 51.43 | -0.11 | *T. angustifolia* | Yes |
| CYTZ_4ATD | -6.79 | 39.21 | *T. domingensis* | Yes |
| E74B | 45.03 | -63.50 | *T. angustifolia* | Yes |
| EDD_44_1 | 47.37 | -68.33 | *T. latifolia* | No |
| EDW_2_15 | 47.37 | -68.33 | *T. latifolia* | No |
| EDW_3_6 | 47.37 | -68.33 | *T. latifolia* | No |
| EL14_TA | 52.44 | 5.85 | *T. latifolia* | No |
| ELTA_01 | 52.44 | 5.85 | *T. angustifolia* | No |
| ELTA_08 | 52.44 | 5.85 | *T. angustifolia* | No |
| FRD47_2 | 45.96 | -66.64 | *T. latifolia* | No |
| FRTD_1 | 41.64 | 13.34 | *T. domingensis* | Yes |
| GORE_3 | 43.68 | -70.44 | *T. latifolia* | No |
| HAW_03_12 | 44.60 | -63.55 | *T. latifolia* | No |
| HAW_10_9 | 44.60 | -63.55 | *T. latifolia* | No |
| HM3TL | 47.51 | 19.04 | *T. latifolia* | No |
| HOR_6 | 48.14 | 26.51 | *T. latifolia* | No |
| HUF_13TA | 47.69 | 17.65 | *T. angustifolia* | No |
| HUS_TL1 | 47.51 | 19.04 | *T. latifolia* | No |
| IP2 | 44.46 | -64.32 | *T. latifolia* | No |
| IR_36 | 44.97 | -64.06 | *T. latifolia* | No |
| IR_39 | 44.97 | -64.06 | *T. latifolia* | No |
| IR_41 | 44.97 | -64.06 | *T. latifolia* | No |
| KL14_TL | 46.64 | 14.31 | *T. latifolia* | No |
| KL1TL | 46.64 | 14.31 | *T. latifolia* | No |
| LIETD_01 | 42.21 | 2.61 | *T. domingensis* | Yes |
| LIETD_12 | 42.21 | 2.61 | *T. domingensis* | Yes |
| LiETL_11 | 42.21 | 2.61 | *T. latifolia* | No |
| M23 | 45.08 | -64.49 | *T. latifolia* | No |
| M30 | 45.08 | -64.49 | *T. latifolia* | No |
| M37 | 45.08 | -64.49 | *T. latifolia* | No |
| M85 | 45.08 | -64.49 | *T. latifolia* | No |
| MBNK_2_8 | 51.51 | -97.00 | *T. latifolia* | No |
| MOD_01_0 | 46.98 | -70.55 | *T. latifolia* | No |
| MOD_02_1 | 46.98 | -70.55 | *T. latifolia* | No |
| MOD_04_1 | 46.98 | -70.55 | *T. latifolia* | No |
| MOD_05_2 | 46.98 | -70.55 | *T. latifolia* | No |
| MOD_07_0 | 46.98 | -70.55 | *T. angustifolia* | No |
| MOD_08_1 | 46.98 | -70.55 | *T. latifolia* | No |
| MOD_14_0 | 46.98 | -70.55 | *T. latifolia* | No |
| MOD_16_1 | 46.98 | -70.55 | *T. latifolia* | No |
| MOD_17_1 | 46.98 | -70.55 | *T. latifolia* | No |
| MOD_18_0 | 46.98 | -70.55 | *T. latifolia* | No |
| MOD_25_2 | 46.98 | -70.55 | *T. latifolia* | No |
| MOD_26_2 | 46.98 | -70.55 | *T. latifolia* | No |
| MOD_29_1 | 46.98 | -70.55 | *T. latifolia* | No |
| MOD_35_3 | 46.98 | -70.55 | *T. latifolia* | No |
| MOD_45_2 | 46.98 | -70.55 | *T. latifolia* | No |
| MOD_54_0 | 46.98 | -70.55 | *T. latifolia* | No |
| NBU_6 | 42.53 | 2.83 | *T. domingensis* | Yes |
| OR_TA4 | 43.28 | -2.13 | *T. domingensis* | Yes |
| OR_TL5 | 43.28 | -2.13 | *T. latifolia* | No |
| P6 | 45.41 | -64.33 | *T. latifolia* | No |
| P9 | 45.41 | -64.33 | *T. latifolia* | No |
| PA2 | 33.19 | -112.15 | *T. domingensis* | No |
| PIW_04_6 | 43.84 | -79.10 | *T. angustifolia* | No |
| PIW_09_1 | 43.84 | -79.10 | *T. angustifolia* | No |
| PIW_13_0 | 43.84 | -79.10 | *T. angustifolia* | No |
| PIW_16_2 | 43.84 | -79.10 | *T. angustifolia* | No |
| PIW_24_3 | 43.84 | -79.10 | *T. angustifolia* | Yes |
| PIW_25_3 | 43.84 | -79.10 | *T. angustifolia* | Yes |
| PWE19TL | 51.62 | -3.94 | *T. latifolia* | No |
| RGT2 | 45.48 | -74.30 | *T. angustifolia* | No |
| SAD_17_3 | 45.27 | -66.09 | *T. latifolia* | No |
| SP2_TA | 52.41 | 4.68 | *T. angustifolia* | No |
| TAKT_1 | 45.27 | 37.38 | *T. angustifolia* | No |
| TAKT_2 | 45.27 | 37.38 | *T. angustifolia* | Yes |
| TALD_1 | 55.73 | 37.61 | *T. angustifolia* | Yes |
| TAVG_1 | 55.32 | 40.37 | *T. angustifolia* | Yes |
| TAVG_2 | 55.32 | 40.37 | *T. angustifolia* | Yes |
| TA_01_F | 26.56 | -81.95 | *T. latifolia* | No |
| TB7 | 44.39 | -64.25 | *T. latifolia* | No |
| TDD_04 | 43.09 | 47.46 | *T. domingensis* | Yes |
| TDD_08 | 43.09 | 47.46 | *T. domingensis* | Yes |
| TDD_09 | 43.09 | 47.46 | *T. domingensis* | Yes |
| TDD_15 | 43.09 | 47.46 | *T. domingensis* | Yes |
| TDD_16 | 43.09 | 47.46 | *T. latifolia* | No |
| TDD_17 | 43.09 | 47.46 | *T. latifolia* | No |
| TDD_18 | 43.09 | 47.46 | *T. domingensis* | No |
| TDD_22 | 43.09 | 47.46 | *T. domingensis* | No |
| TDD_30 | 43.09 | 47.46 | *T. domingensis* | No |
| TDD_32 | 43.09 | 47.46 | *T. domingensis* | Yes |
| TDD_33 | 43.09 | 47.46 | *T. domingensis* | Yes |
| TDD_35 | 43.09 | 47.46 | *T. latifolia* | No |
| TDD_36 | 43.09 | 47.46 | *T. latifolia* | Yes |
| TDD_37 | 43.09 | 47.46 | *T. latifolia* | No |
| TDD_39 | 43.09 | 47.46 | *T. latifolia* | No |
| TDD_40 | 43.09 | 47.46 | *T. domingensis* | No |
| TDD_41 | 43.09 | 47.46 | *T. latifolia* | No |
| TDD_42 | 43.09 | 47.46 | *T. latifolia* | No |
| TDD_43 | 43.09 | 47.46 | *T. angustifolia* | Yes |
| TDD_44 | 43.09 | 47.46 | *T. domingensis* | Yes |
| TDD_46 | 43.09 | 47.46 | *T. domingensis* | No |
| TDD_47 | 43.09 | 47.46 | *T. domingensis* | No |
| TDD_49 | 43.09 | 47.46 | *T. angustifolia* | Yes |
| TDD_50 | 43.09 | 47.46 | *T. domingensis* | No |
| TDD_51 | 43.09 | 47.46 | *T. domingensis* | Yes |
| TDD_52 | 43.09 | 47.46 | *T. domingensis* | Yes |
| TDD_53 | 43.09 | 47.46 | *T. domingensis* | No |
| TDD_54 | 43.09 | 47.46 | *T. domingensis* | Yes |
| TLIK_01 | 45.09 | 147.89 | *T. latifolia* | No |
| TLIK_03 | 45.09 | 147.89 | *T. latifolia* | No |
| TLIK_06 | 45.09 | 147.89 | *T. latifolia* | No |
| TLIK_15 | 45.09 | 147.89 | *T. latifolia* | No |
| TLIK_24 | 45.09 | 147.89 | *T. latifolia* | No |
| TLMK_1 | 45.21 | 38.89 | *T. latifolia* | No |
| TLPK_1 | 45.21 | 38.89 | *T. latifolia* | No |
| TLSK_02 | 47.06 | 142.53 | *T. latifolia* | No |
| TLSK_04 | 47.06 | 142.53 | *T. latifolia* | No |
| TLSK_11 | 47.06 | 142.53 | *T. latifolia* | Yes |
| TLSK_12 | 47.06 | 142.53 | *T. latifolia* | No |
| TMO2_7 | 46.09 | -64.78 | *T. latifolia* | No |
| TXKT_1 | 45.29 | 37.41 | *T. angustifolia* | Yes |
| TXKT_2 | 45.29 | 37.41 | *T. angustifolia* | Yes |
| TXKT_3 | 45.29 | 37.41 | *T. angustifolia* | Yes |
| TXKT_4 | 45.29 | 37.41 | *T. angustifolia* | Yes |
| TXKT_6 | 45.29 | 37.41 | *T. angustifolia* | Yes |
| TXVG_1 | 55.32 | 40.37 | *T. angustifolia* | Yes |
| VFW6 | 49.63 | -125.40 | *T. latifolia* | No |
| Vikram_25 | 44.30 | -78.32 | *T. angustifolia* | Yes |
| Vikram_33 | 44.30 | -78.32 | *T. angustifolia* | No |
| Vikram_76 | 44.30 | -78.32 | *T. latifolia* | No |
| Vikram_94 | 44.30 | -78.32 | *T. angustifolia* | Yes |
| WBD20_1 | 44.28 | -84.23 | *T. angustifolia* | No |
| WBD_27_4 | 44.28 | -84.23 | *T. angustifolia* | No |
| WBD_28_3 | 44.28 | -84.23 | *T. angustifolia* | No |
| WBD_32_1 | 44.28 | -84.23 | *T. angustifolia* | Yes |
| WBD_33_1 | 44.28 | -84.23 | *T. angustifolia* | No |
| WBW5_4 | 44.28 | -84.23 | *T. latifolia* | No |

## Notes

### Competing Interest Statement

The authors have declared no competing interest.

### Summary of Updates

We have made some adjustments in terms of the paper's wording and structure, which are aimed at further enhancing the readability and flow of the document. These changes solely pertain to the paper's form and do not impact its content or the core findings.

